# Assessment of a high-throughput mass spectrometry method to accelerate biomarker discovery in clinical cancer cohorts using volumetric absorptive microsampling (VAMS) devices

**DOI:** 10.1101/2024.10.10.617693

**Authors:** N. Lucas, C. Hill, D. Pascovici, R. McMahon, B. Herbert, E. Karsten

## Abstract

Identification of biomarkers of early-stage disease typically requires analysis of very large cohorts which can only be reasonably achieved using high-throughput methods. We have developed and optimised novel methods for whole blood analysis using volumetric absorptive microsampling devices to produce over 3000 protein identifications by LCMS on a Q Exactive HF-X Orbitrap. These methods were tested using a set of whole blood samples from lung cancer patients and matching healthy controls finding 455 differentially expressed proteins, using a mid-throughput method enabling analysis of 18 samples per day. To increase throughput for larger clinical cohorts, a 60-sample per day method was tested on a Sciex ZenoTOF 7600. The high-throughput method produced 1.5-fold fewer protein identifications and a higher overall % CV compared to the mid-throughput method. Despite the lower numbers, it produced a set of 36 disease-relevant and discriminatory differentially expressed proteins that, using a machine learning model, could differentiate between the disease and control samples with an area under the ROC curve (AUC) of 88.9% using random forest algorithms. These data support the use of high-throughput mass spectrometry methods to screen large cohorts for diagnostic biomarkers that can then be followed up with more targeted analyses.

## Introduction

Large scale biomarker discovery in biological fluids is being enabled by the development of technologies that quantify thousands of proteins in small volumes, typically less than 10uL. Immunoassay platforms such as Olink and SomaScan now survey panels of thousands of proteins in runtimes measured in hours rather than days^1, 2^. Mass spectrometry (MS) instruments are now sensitive enough to quantify thousands of proteins from a single cell or microlitre volumes of fluid such as plasma in runtimes of less than 20 minutes^3–5^. Unfortunately, when it comes to MS analysis of blood the issue of dynamic range has still been a major pitfall for large scale proteomics^6^. To achieve protein identification numbers greater than 2,000 usually requires depletion and/or extensive fractionation^7, 8^ of the sample. More recently there has been shift in enrichment techniques for plasma proteomics with the development of nanoparticle bead enrichment strategies, which decrease the dynamic range through competitive binding. While these techniques can reduce the dynamic range and enable low abundance protein detection, they are usually labour intensive, slow and expensive^3, 9, 10^.

Most proteomics studies that focus on blood use plasma or serum prepared from a venipuncture, which requires specialist collection by a phlebotomist. However, a growing interest in remote and at-home sampling has led to publications using microsampling to collect whole blood, commonly in the form of a finger-prick dried blood spot^11^. These samples can be easily self-collected by the donor, which eliminates the need for specialist collection and enables patient-centric longitudinal collection protocols. Conventional dried blood spots based on filter paper have several advantages over traditional blood collection, including their ease of collection, transport and storage. They do suffer from known disadvantages primarily due to the varying concentration of red blood cells in patients’ blood, known as haematocrit effect. High concentrations of red blood cells result in less diffusion through the paper matrix and consequently a smaller more concentrated spot^12^. One solution to this issue was the development of volumetric absorptive microsampling devices (VAMS)^13^ which collect a specific volume of blood, independent of the haematocrit, onto a hydrophilic polymeric tip attached to a spindle. These devices have been designed to function in a standard 96-well format, which is compatible with widely used laboratory systems and enables the development of automated preparation methods^14–16^.

An important step in optimising methods for clinical specimen analysis is increasing sample throughput. Immunoassay platforms now offer large, multiplexed protein panels, with automated sample processing capable very high throughput of samples per day (SPD). Olink and SomaScan for example report the capacity to run between 120 – 1000 SPD depending on the assay. The disadvantage of these assays however is that they require antibodies or aptamers that are specific to sets of known proteins only. Comparatively, MS is exploratory which can aid in the identification of novel biomarkers and is applicable to a wider range of samples. However, because the peptide mixture is separated by liquid chromatography before MS detection there is usually a much lower throughput capacity (< 50 SPD). In order for LC-MS to be competitive with immunoassay technologies, it is necessary to explore methods of increasing the throughput of samples.

Following on from an earlier publication^17^, we have further optimised a method for whole blood analysis using MS which enables quantification of more than 3,000 proteins from VAMS. Using VAMS devices, we found that washing dried blood samples with various normal salt solutions removes a significant proportion of haemoglobin, albumin and other highly abundant proteins, decreasing dynamic range of the sample. This method has advantages over other enrichment, depletion, or fractionation methods as it requires minimal sample handling after collection. In this paper, we have applied this method to a clinical cohort of 34 samples (18 control and 16 lung cancer patients) and investigated the use of both mid- (18 samples per day (SPD)) and high-throughput MS (60 SPD) methods on the quantity and quality of biomarkers as well as the benefits for large scale clinical blood proteomics.

## Methods

### Blood collection

Frozen venous whole blood cell pellets from 16 patients with non-small cell lung cancer (NSCLC) and 18 age and sex matched controls were acquired from Precision Med, Inc (Solana Beach, CA, USA). Informed consent was obtained from all subjects involved in the study.

### Sample processing

Biospecimen were randomised into 2 processing batches. In these batches, the samples were thawed and diluted in an equal volume of PBS. A Neoteryx Mitra® microsampling tip (30 µL) was then dipped into the diluted whole blood cell pellets until the tip had wicked up the blood and was full. A single 30 µL tip was prepared for each sample. All samples were then air-dried for 5 minutes at room temperature before being transferred to a foil ziplock bag with desiccant. These samples were left to completely dry for 24 hours at room temperature.

After the 24 hour period, the samples were extracted immediately. The VAMS tips were removed from the plastic spindles and were suspended in 1 mL extraction solution (500 mM lithium chloride + 100 mM Tris) in Eppendorf tubes to incubate at room temperature overnight for 24 hrs whilst gently shaking. Following incubation, the tips were washed twice in extraction solution. The proteins remaining in the tip were digested by suspending the washed tips in 100 µL lysis buffer (1 % sodium-deoxycholate (w/v), 10 mM TCEP (tris(2-carboxyethyl)phosphine, 40 mM chloroacetamide, 100 mM triethylammonium bicarbonate buffer, pH 8) and were heated at 95 °C for 10 mins with agitation (Thermomixer, Eppendorf). After heating, 1 µg trypsin was added (1 µg/µL in lysis buffer) and the sample was incubated overnight at 37 °C (19 hours). Samples were then diluted with 800 µL of 0.5 % formic acid (FA) and the tip was removed from the liquid sample and discarded. The remaining sample was centrifuged (16,000 *g*, 10 mins) and the supernatant was transferred to a conditioned solid phase extraction cartridge (Oasis HLB 1cc cartridge, 10 mg sorbent, 30 µm). Sample was desalted using solid phase extraction. Peptide concentration was determined using Nanodrop spectrophotometer (Thermo Fisher). Samples were frozen (−80 °C) until analysis.

### Sample analysis

All samples were run on 2 instruments, to compare a mid-throughput and high-throughput method. For both instruments, a HeLa digest was run in triplicate and used to benchmark both instruments.

**Figure 1.**
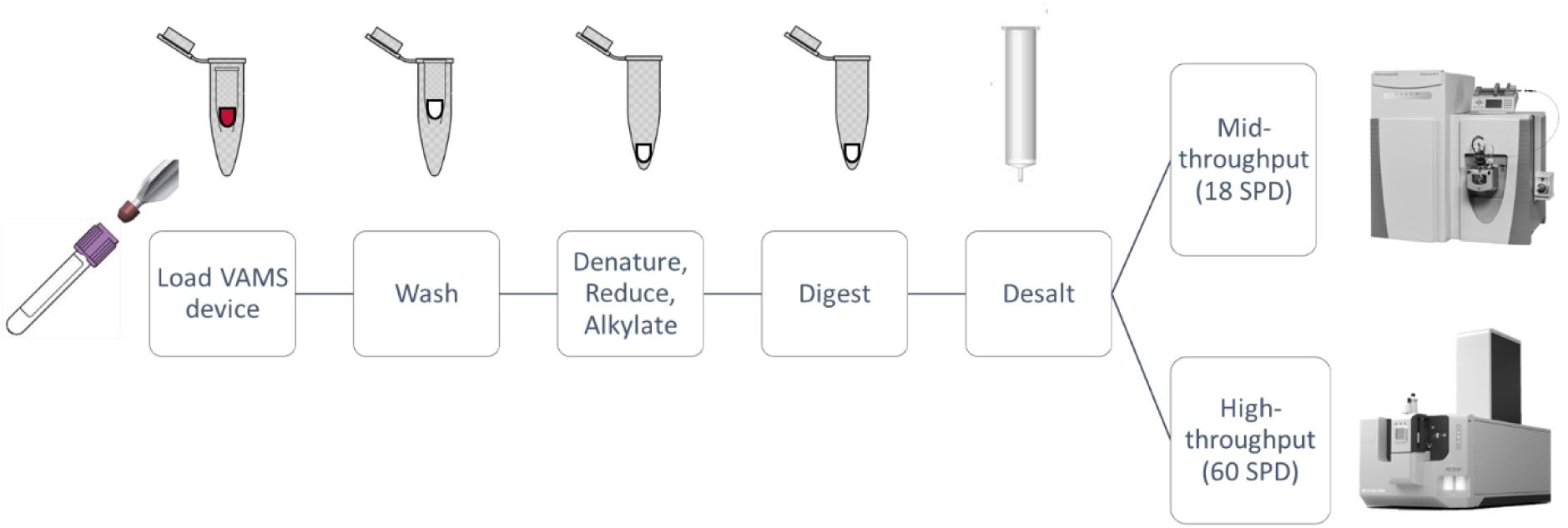
Schematic of sample preparation analysis methods.

#### Mid-throughput method (18 samples/day) - Thermo Q Exactive H-FX

Peptides (0.5 µg) were loaded directly onto a nano LC column (75 µm × 50 cm 1.9 µm C18 ReproSil-Pur 120 C18-AQ) in 97 % solvent A (0.1 % FA) and 3 % solvent B (80 % acetonitrile, 0.1 % FA) at 500 µL/min for 12 mins using an Ultimate 3000 UHPLC (Thermo Scientific, CA, USA). The flow rate was decreased to 300 µL/min and peptides were eluted over 62.5 min gradient to a maximum of 30 % solvent B. The gradient was then stepped up to 60 % solvent B over 2.5 min, before washing in 98 % solvent B for 5 mins.

Peptides were detected using a Q Exactive HF-X Orbitrap mass spectrometer (Thermo Scientific, CA, USA). Data was acquired in DIA-mode using the following instrument parameters: Full MS scan resolution 120 K, AGC 3 × 10^6^, maximum injection time 50 ms, scan range 350-1650 m/z, followed by 20 DIA variable windows to scan 350-1650 m/z as per **Table S1** with settings of Resolution 30 K, AGC 3 × 10^6^, maximum injection time set to auto. The collision energy was stepped at 22.5, 25, 27.5, default state was 3.

#### High-throughput method (60 samples/day) - ABSciex ZenoTOF 7600

Peptides (0.5ug) were loaded onto Evotip Pure tips (EV-2011) according to manufacturer’s instructions. Sample containing tips were then loaded onto Evosep One (EV-1000) LC system (Evosep, Denmark) coupled to ZenoTOF 7600 mass spectrometer (SCIEX, MA, USA). Standard Evosep 60 samples per day (SPD) method was used with Evosep column EV-1109 analytical column (Dr Maisch C18 AQ, 1.5um beads, 150um ID, 8cm long). Data was acquired using Zeno SWATH with the following instrument parameters: Optiflow 1-50uL microflow sources set to 4500 V ion spray voltage; curtain gas at 25; temperature 150°C; source gas 1 set to 12; and source gas 2, 60; and column oven set to 40°C. For DIA/SWATH acquisition peptide spectra were acquired with the LC MS/MS using 67 variable windows as shown in **Table S2**. TOF MS scan range 450 - 850 m/z with a 1 Da window overlap; declustering potential (DP) was set to 80; DP spread 0; collision energy (CE) is as shown in **Table S2**, and CE spread was 0. MS2 spectra were collected in the range of m/z 100 to 1800 for 11 ms.

### Data processing

All files were processed through DIA-NN 1.8.1 (Data-Independent Acquisition by Neural Networks) as two independent searches for either the mid- or high-throughput method. Global settings are as follows: Output was filtered at 0.01 FDR; Deep learning was used to generate a new in silico spectral library from peptides provided; Library-free search enabled; N-terminal methionine excision enabled; In silico digest involved cuts at K*,R*; Maximum number of missed cleavages set to 1; Min peptide length set to 7; Max peptide length set to 30; Min precursor charge set to 2; Max precursor charge set to 5; Cysteine carbamidomethylation enabled as a fixed modification; A spectral library was created from the DIA runs and used to reanalyse them. DIA-NN optimised the mass accuracy automatically using the first run in the experiment. For the mid-throughput run the following settings were used Min fragment m/z set to 350; Max fragment m/z set to 1650; Min precursor m/z set to 100; Max precursor m/z set to 2000. For the high-throughput run the following settings were used Min fragment m/z set to 450; Max fragment m/z set to 850; Min precursor m/z set to 100; Max precursor m/z set to 1800.

### Statistical analysis

Statistical analysis was performed on DIA-NN output files (version. 1.8.1). Files were analysed using R Studio and Mass Dynamics (accessed 24-Jan-2024). Mass spectrometry data was filtered to remove sparse proteins, with fewer than 3 quantitated values in both groups. Remaining protein quantitation was zero or missing values were imputed using the default Perseus-style imputation. Median normalisation was carried out before data imputation and no batch normalisation was undertaken. After data imputation, differential expression analysis was carried out. Initially the data was compared between groups with a two-sample t-test and proteins were regarded as differentially expressed if the false discovery rate corrected p-value was less than 0.5. Graphing was performed with GraphPad Prism (vers. 10.1.2), R-Studio, and Mass Dynamics (accessed 24-Jan-2024). Machine learning based modelling was performed using ProMor package for R Studio^18^. Methods followed are as published. Briefly, log2 transformed DIA-NN protein data matrix were uploaded and proteins with high levels of missing data filtered out (>34% missing values in at least one of the groups). Data was then imputed (‘minProb’ method) and normalised (‘quantile’ method) and after a differential expression analysis (log2 fold change >1 at an adjusted P-value <0.05) a model data frame was created using default settings. Data was split into a training set (28 samples) and a test set (6 samples) for each mid and high throughput methods. Proteins from this model were used with machine learning algorithms k-nearest neighbours (knn) and random forest (rf) to produce AUC curves.

## Results and Discussion

### Overall numbers of proteins

All samples were analysed using both a mid- and high-throughput method, and the results of each instrument were compared. The mid-throughput method was run on a Q Exactive HF-X mass spectrometer (Thermo Scientific) using a 70-minute gradient, with an overall capacity of 18 samples per day. The high-throughput method was run on an ZenoTOF 7600 mass spectrometer (Sciex) using 24-minute gradient, with an overall capacity of 60 samples per day. The high-throughput method was 3.3-fold faster than the mid-throughput method.

Due to the longer gradient and LC column of the mid-throughput method, significantly more proteins were identified in comparison to the high-throughput method, with a mean and SD of 3370 ± 117 and 1483 ± 137 for each respectively (**Figure 2a**). Per minute of run time however, the high-throughput produced 1.5-fold more IDs (**Figure 2b**). These trends were consistent across all samples, including when stratified by disease status (**Figure 2c-d**).

**Figure 2.**
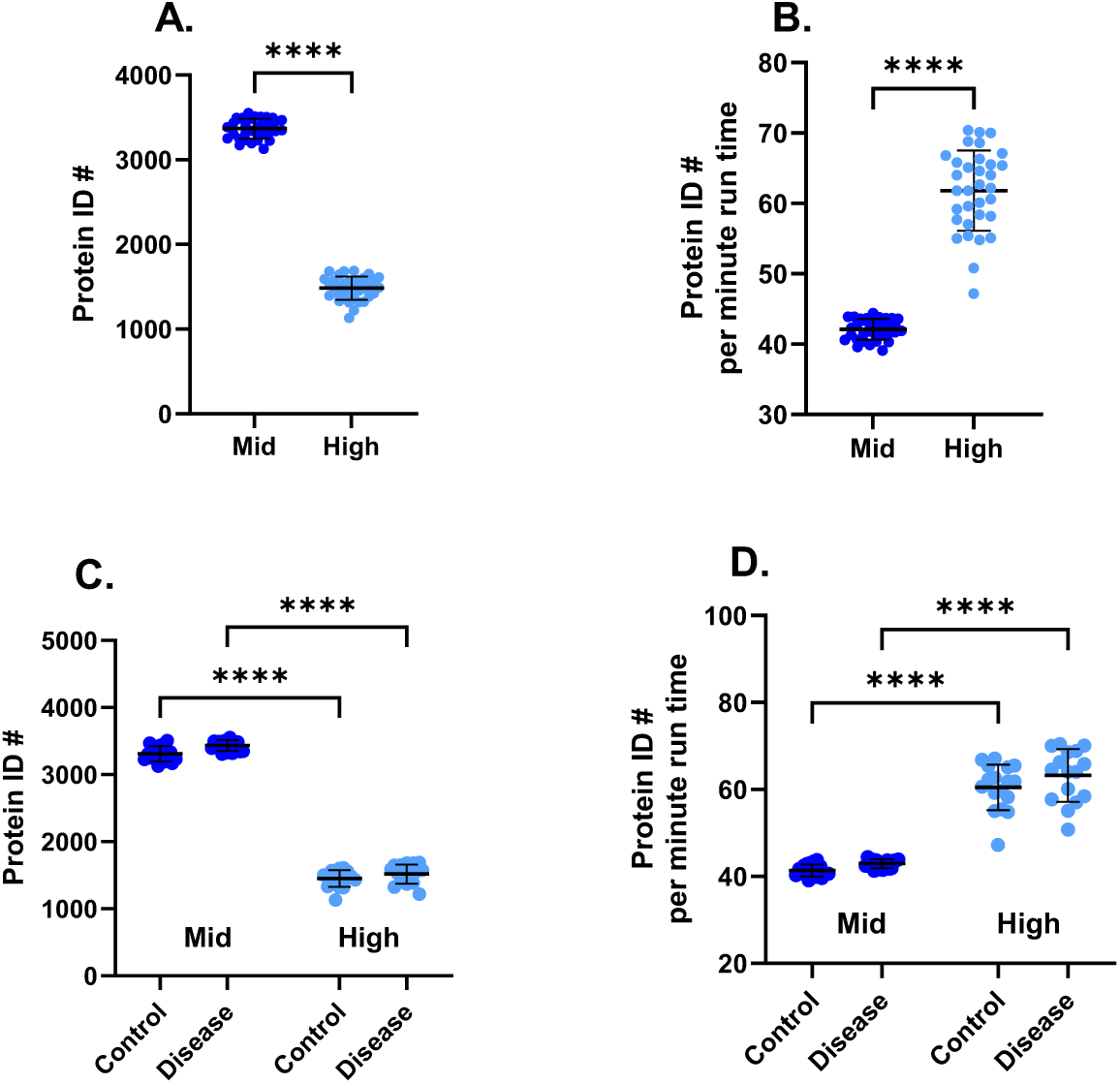
Number of protein IDs for samples analysed with a mid-throughput method (compared to a high-throughput method for (A) total acquisition time, (B) per minute of acquisition time, (C) total acquisition time stratified by disease group, or (D) per minute of acquisition time stratified by disease group. Data are mean ± SD, data is significantly different (*) if p < 0.05.

It is important to note that both methods performed admirably given the sample analysed. Mc Ardle et al. analysed whole blood VAMS using mid- (20 SPD) and high- (57 SPD) throughput method and reported a similar scale reduction with the high-throughput method of 1.5-fold, but were only able to identify 518 and 350 proteins with each method respectively^16^. This works out to be 6.7 and 4.2-fold fewer protein identifications for each method compared to our results.

Our overall higher numbers of identifications in this study are likely due to our novel sample processing methods. This method was first described in 2022^17^ and has since been optimised as outlined here. Although the details of sample extraction were not included in Mc Ardle et al^16^, they referenced the method outlined in van den Broek et al^14^ which describes a standard extraction and digestion protocol for VAMS. Briefly, this includes an incubation in an extraction solution then all subsequent sample processing is performed in a single tube containing both the resulting liquid sample and extracted tip (both of which is heavily contaminated with hemoglobin and other high abundance proteins). Our methods comparatively utilise an initial extraction that is discarded and all additional sample processing is performed on the sample remaining in the tip (**Figure 1**). This process enables removal of high abundance proteins and reduces the dynamic range of the resulting sample.

### Analysis Cost

An important factor in large biomarker studies is the cost of preparing and acquiring sample data. We did a basic cost analysis using commercial prices per hour provided by a mass spectrometry analysis facility (based in Australia) **Figure 3**. Although the consumables for the high-throughput method are more expensive per unit, the shorter run time makes the total cost of this method approximately one-third the cost of the mid-throughput method (**Figure 3**). When calculated per analyte detected, the costs for both methods are comparable at approximately $AUD 0.02 per analyte. Notably, the costs for each method are 10-fold more affordable than the discovery scale immunoassay equivalents, such as SomaScan (up to 7000 analytes) or Olink Explore (up to 3000 analytes) which both cost approximately $AUD 0.20 per analyte.

**Figure 3.**
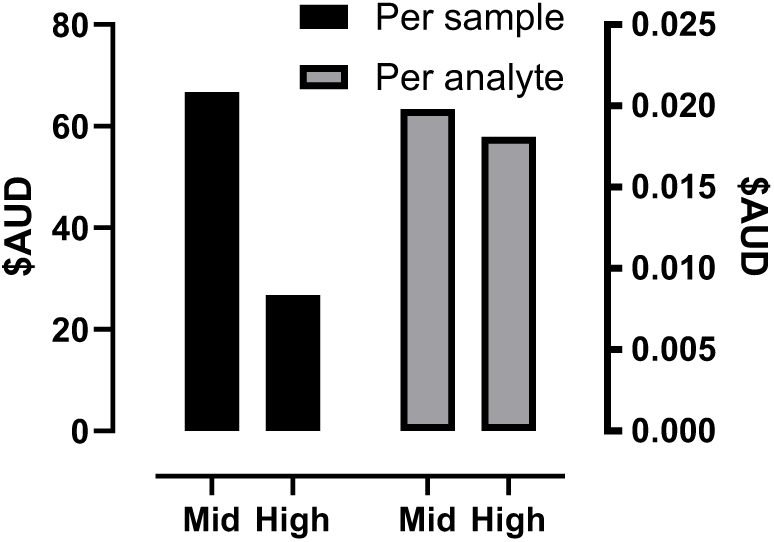
Cost comparison in $AUD between mid- and high-throughput proteomic analysis per sample (left y-axis) and per analyte per sample (right y-axis).

### Data Variability

The protein intensity data before and after imputation was found to be more variable for the high-throughput method, as illustrated by the increased median and spread of the % CVs (**Figure 4a**). The trend is again consistent across both the control samples and the disease samples (**Figure 4b-c**), with slightly improved correlation between methods for the disease samples (**Figure 4e**). Benchmarking for both methods was done by running HeLa cell lysate in triplicate, and median % CVs were below 10 % for each method (Supplementary Figure S1-A). After reviewing peak widths and scans post-acquisition there was more variation found in the high-throughput method (Supplementary Figure S2 (B-C)), not only between mid- and high-throughput methods but also between the clinical samples and the HeLa. This was not as evident using the mid-throughput method. This is most likely due to dynamic range issues when using a shorter gradient, as high abundant proteins can contribute to longitudinal diffusion of peaks and could potentially be further optimised.

**Figure 4.**
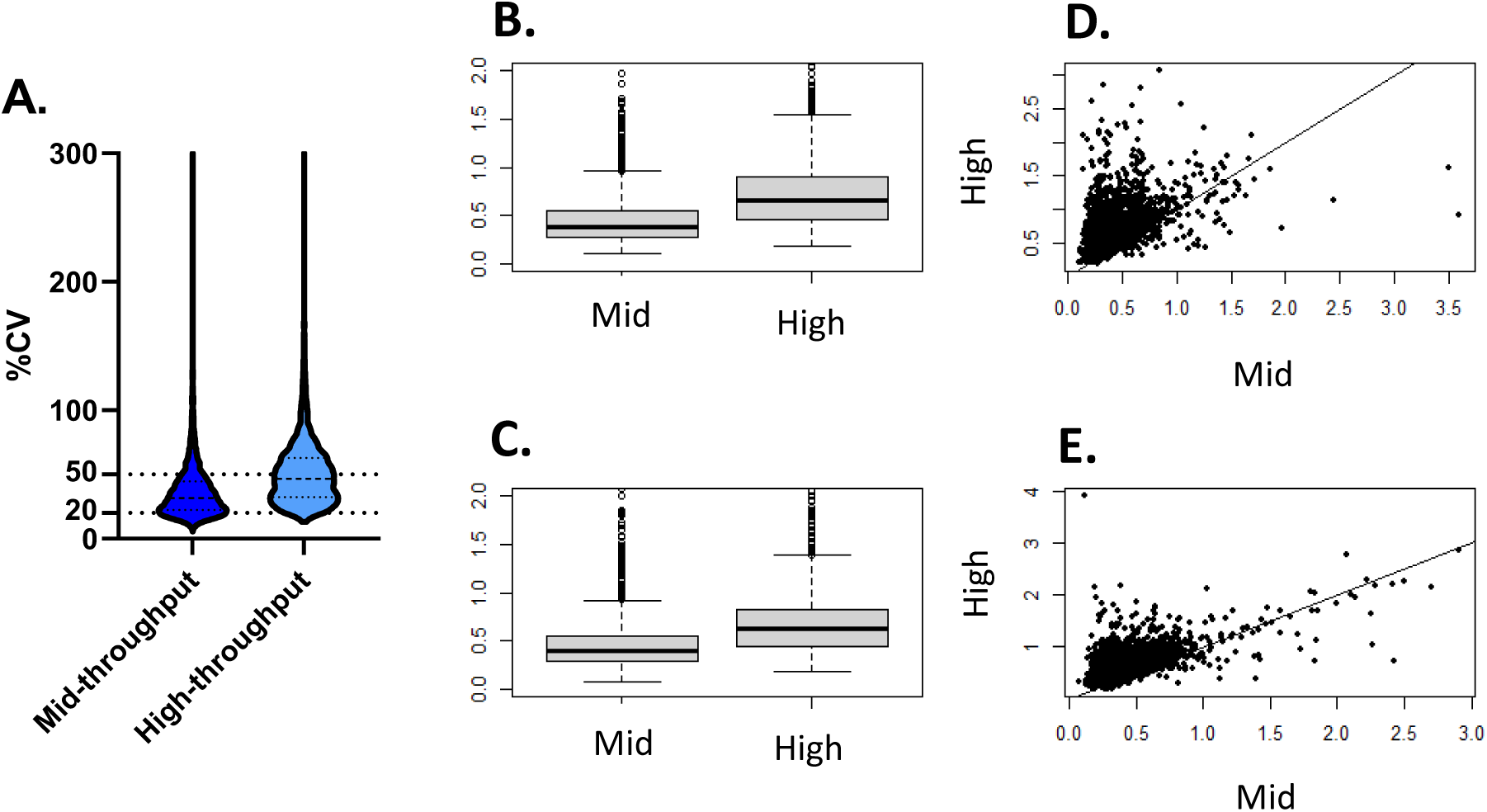
% CV depicted as (A) violin plots of all samples analysed using mid- and high-throughput methods; boxplots of samples stratified by disease status, (B) control and (C) disease; scatter plots comparing CV of mid- and high-throughput methods stratified by (D) control and (C) disease.

### Data completeness of proteins across samples

The number of proteins with quantitation across samples was determined for each method. For the mid-throughput method, more than two thirds of the proteins were quantitated in all samples, while for the high-throughput method, less than 50 % of the data was quantitated in all samples (**Figure 5**) thus demonstrating greater sparsity in the high-throughput dataset. Although variability in protein detection across samples is expected for a clinical cohort, this variability should be reflected with both methods. The higher variability in the high-throughput method is consistent with the more variable % CVs as discussed above. One of the potential implications of the higher variability is the need for more imputation or rejection of data, which will have downstream effects on biomarker discovery. However, the ability to be able to survey more samples in a shorter time-frame will also provide greater statistical power.

**Figure 5.**
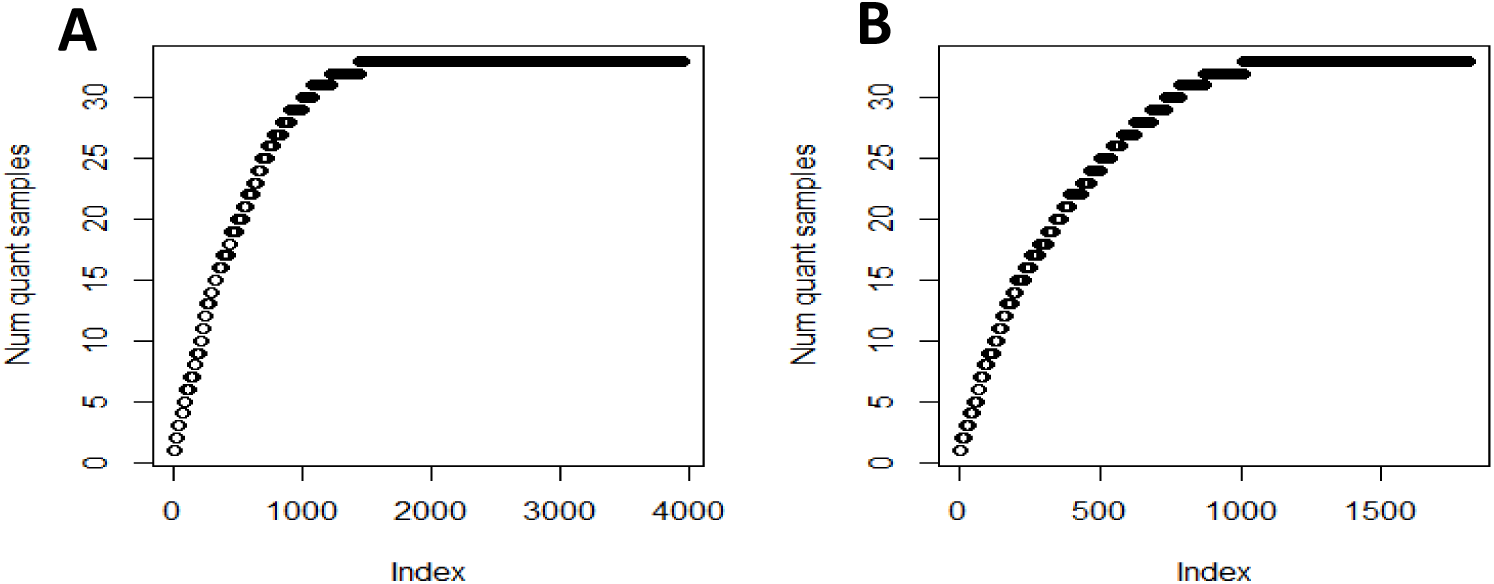
The number of samples quantitated for ranked proteins with the (A) mid- and (B) high-throughput methods.

### Dynamic Range

One of the main challenges with increasing throughput using MS methods, is the need for higher flow rates and smaller length columns. This leads not only to a trade off in protein numbers detected but an increase in dynamic range due to co-eluting peaks caused by peak compression. The dynamic range for each method and instrument was first determined by assaying HeLa cell lysates in triplicate (**Figure 6a**). Detected proteins were ranked by LFQ intensity and means were plotted. Almost double the number of proteins (1.8-fold) were detected using the mid-throughput method for HeLa cells with an order of magnitude increase in sensitivity compared to the high-throughput method. The same trend was observed for the blood samples, however with a slightly higher (2.2-fold) number of proteins detected with the mid-throughput method as expected with the higher dynamic range of the blood samples (**Figure 6b**). The dynamic range calculations highlight the strength of the extraction method which performed well for the blood samples compared to HeLa controls, producing thousands of identifications across 5 logs of intensity, a considerable improvement to standard plasma analysis ^6^.

**Figure 6.**
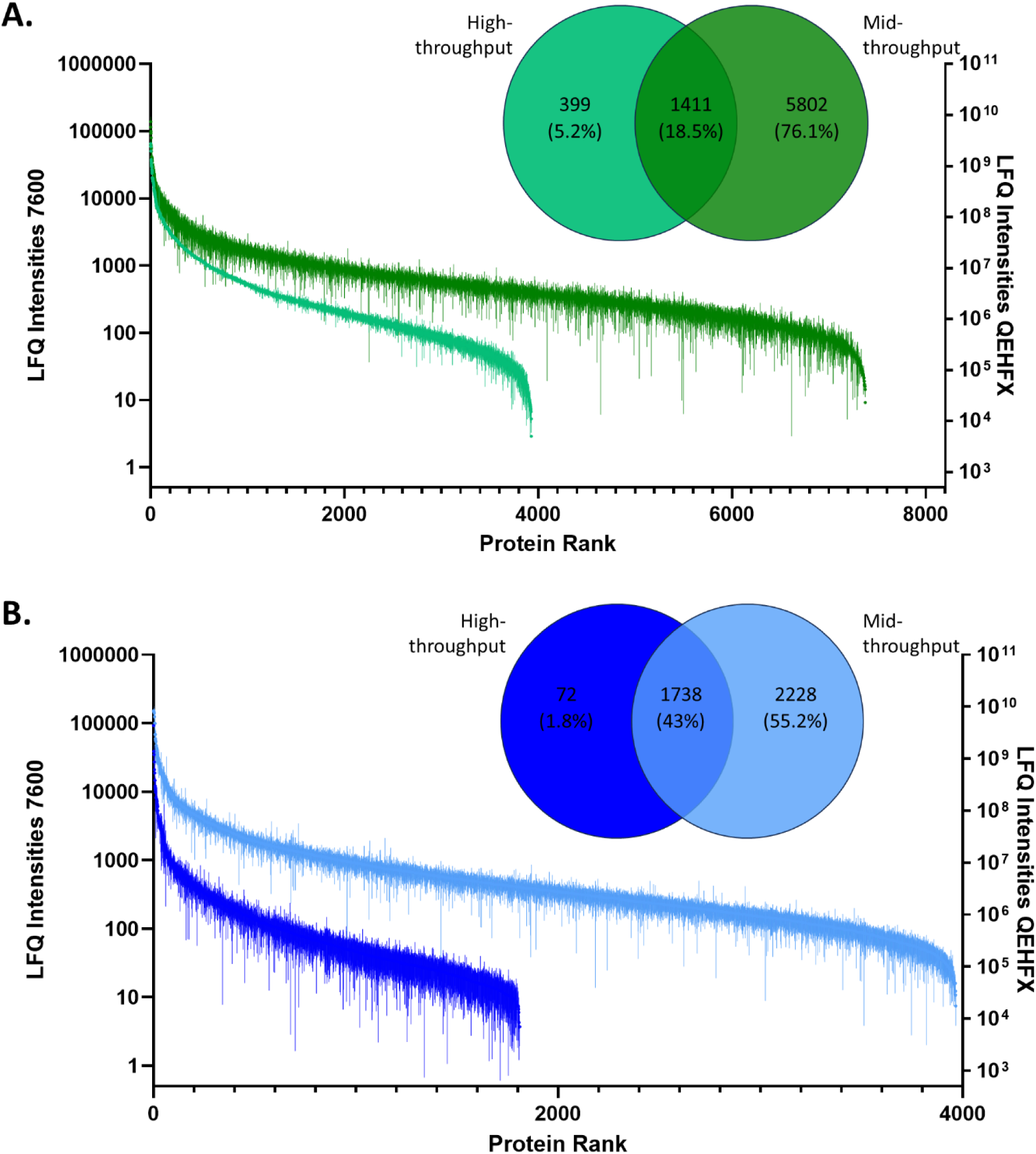
Dynamic range of LFQ intensities of ranked proteins analysed using both high-throughput methods (left y-axis) and mid-throughput methods (right y-axis) for (A) HeLa cell controls (n = 3) and (B) blood samples (n = 34). Data presented as mean ± SD. Overlap in detected proteins illustrated by Venn Diagrams.

### Differentially expressed proteins

Differentially expressed proteins were identified between control and disease samples for both the mid- and high-throughput method. There was good correlation in fold change between protein intensities of control and disease data for both methods (**Figure 7a**). The adjusted *p*-values for the control-disease comparison were found to be overall higher in the high-throughput method (**Figure 7b**), which is consistent with the dataset being more variable (**Figure 4**-**5**).

**Figure 7.**
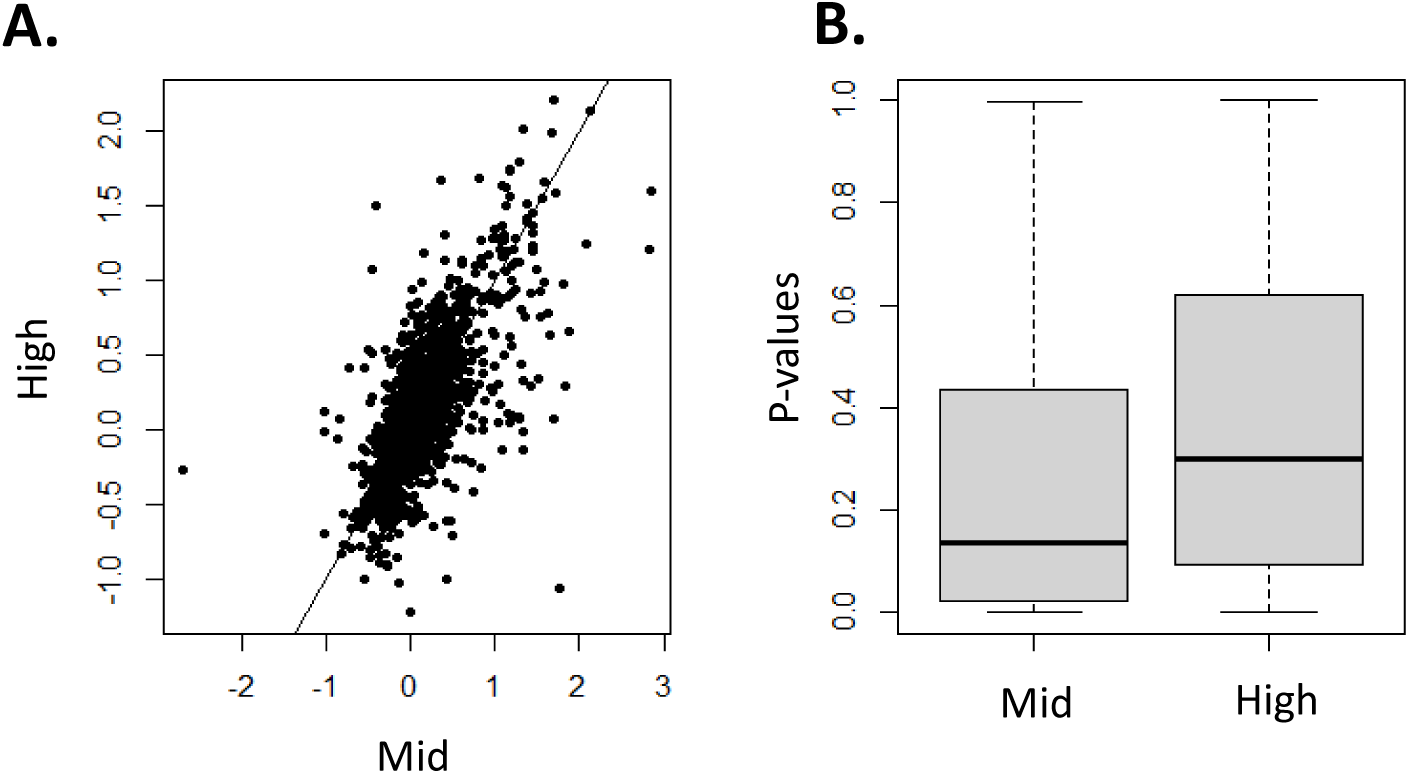
(A) Scatter plot of log2 fold change of protein LFQ intensities between control and disease samples determined by mid-throughput and high-throughput methods and (B) boxplots of unadjusted p-values for differential expression of proteins between control and disease samples using mid- and high-throughput methods.

The distribution of the volcano plots is similar for each method, with a weighting towards up-regulated differentially expressed proteins in the disease samples (**Figure 8a-b**). As anticipated, the high-throughput method had significantly fewer differentially expressed proteins that surpassed the 1.5-fold change threshold and were above the 5% FDR threshold, with a resulting 36 proteins identified compared to the 455 proteins identified in the mid-throughput method (**Figure 8a-b**). Despite the high-throughput method identifying 12.6-fold fewer differentially expressed proteins, 31/36 of these identified proteins overlapped with the mid-throughput results (**Figure 8c-d**). The complete list of these overlapping proteins is included in **Supplementary Table S3**. This demonstrates some utility in the method. The increased speed of the method enables screening of larger sample cohorts, and the overlap in proteins of interest that we observed between the methods demonstrates that not all value is lost by the increase in throughput.

**Figure 8.**
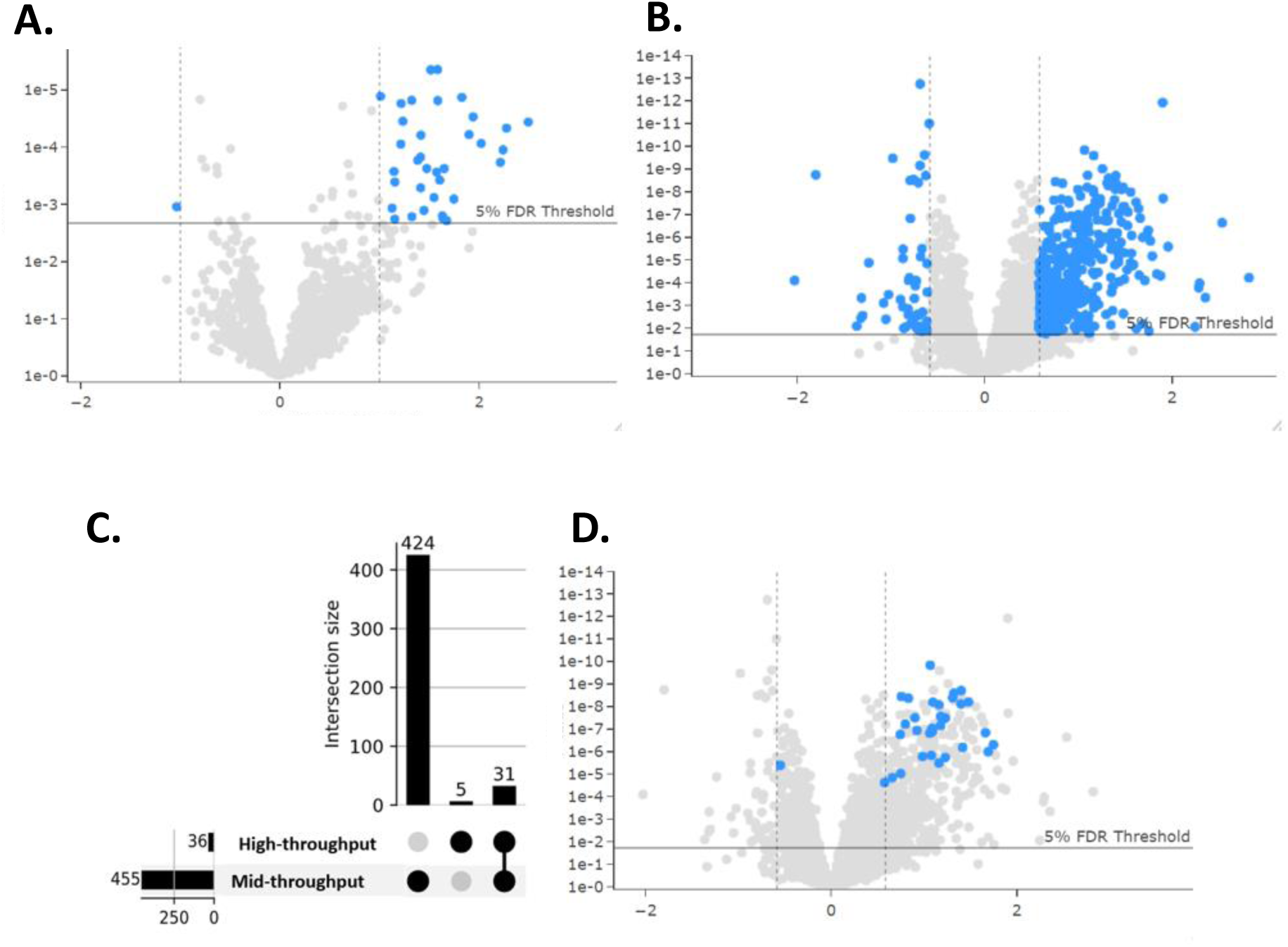
Differentially expressed proteins between controls and disease above the threshold of 5% FDR and fold change ratio of 1.5 (blue dots) as expressed by (A) a volcano plot of results using high-throughput methods, (B) a volcano plot of results using mid-throughput methods, (C) an upset plot of the overlap of proteins identified in each analysis, and (D) a volcano plot of the mid-throughput analysis with the overlapping proteins from both analyses highlighted in blue.

A number of the overlapping proteins have been previously identified as biomarkers of NSCLC. MS analysis of NSCLC tumour tissue identified upregulation of a number of proteins including AZU1, MPO, and LTF^20^, and comprehensive transcriptome analysis identified CYBB and LYZ in stage III tumour tissue^21^, all of which were found using both through-put methods in our study supporting the validity of our results as markers of lung cancer.

### Discrimination of status by machine-learning based modelling

To assess the strength of the differentially expressed proteins identified using the two methods (mid- and high-throughput) machine learning based modelling was performed using the ProMor package for R Studio^18^. A fourteen-marker prediction model was identified for the high-throughput method (A) which gave an area under the ROC Curve (AUC) of 72.2% using k-nearest neighbours (knn) and 88.9% using random forest (rf) algorithms (C). For the mid-throughput method, a six-protein prediction model (B) gave an AUC of 94.4% using k-nearest neighbours and 77.8% using random forest algorithms (D). The analysis demonstrates that even though the high-throughput method identified fewer differential proteins the model proteins still gave acceptable sensitivity and specificity to differentiate cancer patients from healthy controls. The mid-throughput model showed outstanding discrimination using k-nearest neighbours with and AUC of 94.4%, however the sample set is very small (training set of 28 samples and test set of 6) and larger cohorts would be required to validate this further.

There was only one protein common between the mid- and high-throughput models, a serine protease, Myeloblastin (PRTN3). PRTN3 has been linked to KRAS mutations in lung cancer patients^22^. It has also been shown to be linked to poor prognosis in colorectal cancer patients ^23^ and as a potential biomarker for vulvar squamous cell carcinoma patients ^24^. Other proteins in the mid-throughput panel, have also previously been implicated in cancer. Beta-Ala-His dipeptidase (CNDP1) has been shown to be downregulated in NSCLC patients^25^, and used in diagnostic panel^26^. While Omega-amidase NIT2 has been identified as a potential tumour suppressor due to its role in cell growth inhibition ^27^. In the high-throughput panel the two top proteins with the highest importance, intercellular adhesion molecule 3 (ICAM3), myeloid cell nuclear differentiation antigen (MNDA), have both previously been identified as biomarker candidates for diagnostic and therapeutic treatments of lung cancer^28, 29^. ICAM3 has been found to increase cancer cell migration/invasion^28^, whilst MNDA has also been shown to activate immune cells^29^.

**Figure 9.**
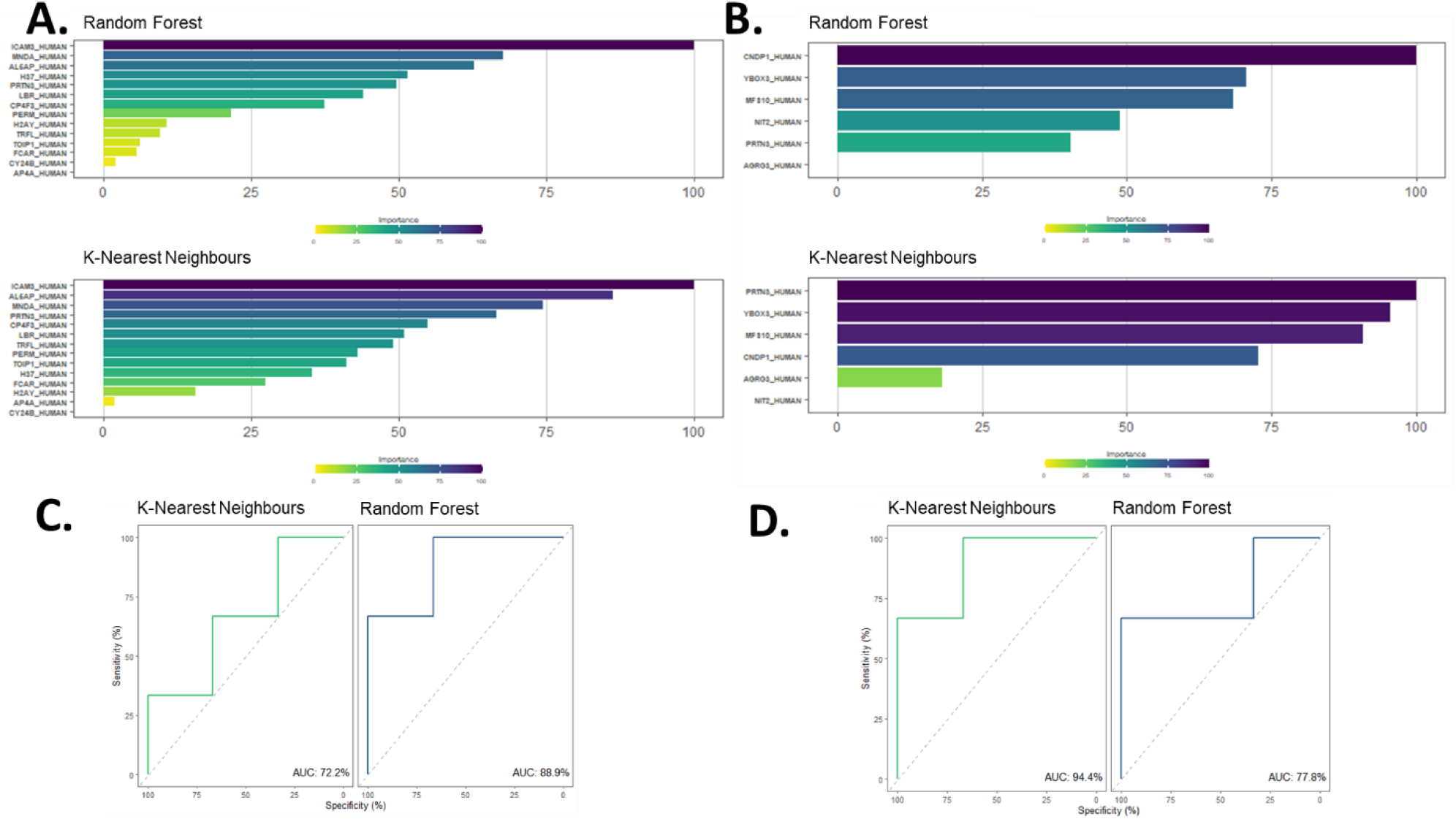
Fourteen-marker prediction model for the high-throughput method (A) Receiver operating characteristic curve (ROC) analysis, using both k-nearest neighbours (knn) and random forest (rf) algorithms (C). Six-protein prediction model for the mid-throughput method (B) and ROC analysis using knn and rf algorithms (D).

### Significant pathways

Reactome ORA pathway analysis was performed on each set of differentially expressed proteins. There were 5 pathways identified that were statistically significant and were common to each method (**Table 1**). All of these pathways are closely related, with neutrophil degranulation, antimicrobial peptides, and ROS and RNS production in phagocytes being pathways under the innate immune system, which is a branch of the immune system, indicating a strong immune system response in the disease cohort that is detected by both methods.

**Table 1.**
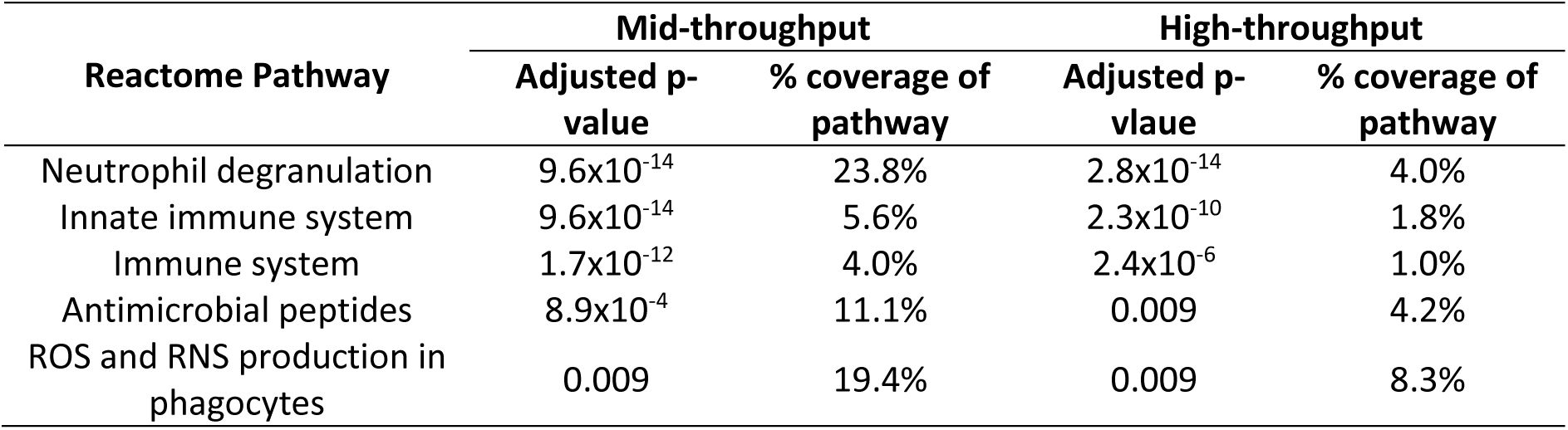
Over-represented reactome pathways identified as statistically significant (with FDR adjustment) for both mid-throughput and high-throughput mass spectrometry methods.

The innate immune system is one of the main pulmonary defences against infection, but in tumours the innate immune system has been found instead to promote disease progression^30^. A 2023 publication of whole blood gene expression in late-stage lung cancer also reported the neutrophil degranulation pathway as highly significant in the lung cancer cohort and found an association with increased neutrophil numbers in whole blood and diagnosis^31^.

The high-throughput method produced one additional statistically significant pathway (interleukin-4 and interleukin-13 signalling) that was not significant with the mid-throughput method, with 4.5% and 4.6% pathway coverage respectively. The mid-throughput method produced an additional 23 pathways that were not identified with the high-throughput method. Of those, four were recognised disease pathways (1. Defective CFTR causes cystic fibrosis; 2. ABC transporter disorders; 3. Disorders of transmembrane transporters; 4. Synthesis and processing of ENV and VPU).

## Conclusions

Improving on an earlier published method we show that the washing of VAMS devices enables the reduction of protein dynamic range in clinical blood samples allowing the quantification of more than 3,300 proteins using a mid-throughput (18 SPD) MS method. Applied to a cohort of 34 lung cancer patients and healthy controls we further demonstrate its ability to find potential biomarkers discriminating between disease status.

To increase throughput for future cohorts, and gain more statistical power, we also analysed the same samples using a high-throughput MS method (60 SPD). With an average of 1,483 proteins quantified per sample we identified 36 differentially expressed proteins in the high-throughput method, compared to 455 found with the mid-throughput method. Although we did see reduced protein identifications and increased variability using the faster method (in both %CVs and data missingness) the ability of the high-throughput method to identify relevant biomarkers and discriminate between healthy and disease patients still proved to be sufficient. One of the most important factors in identifying quality biomarkers, suitable for clinical translation, is reproducibility and this is often achieved by using cohorts with large numbers of samples. Using a high throughput method, which can process 60 samples a day, enabled differential expression biomarker data to be generated in less than a third of the time of conventional mid-throughput methods.

The use of machine learning algorithms identified prediction models of 14- and 9-biomarkers for both high- and mid-throughput methods respectively. Of these several markers including PRTN3, CNDP1, NIT2, MNDA, ICAM3 have been identified previously in lung cancer and show promise as future markers to be included in validation studies. Pathway analysis also produced confirmatory results in identifying known lung cancer associated pathways, such as neutrophil degranulation and the innate immune system.

The combination of our novel sample preparation methods with both the mid- and high-throughput protocols produced high quality data with sufficient quantified identifications to enable differential biomarkers to be defined. This study also demonstrates the potential use for microsampling in cancer biomarker discovery which has real-world clinical utility, providing a path to at-home sampling and patient centric monitoring.

## Acknowledgments

Mass spectrometry analysis was performed at Sydney Mass Spectrometry, University of Sydney.

## Author Contributions

<u>Conceptualisation:</u> N.L., E.K.; <u>Methodology:</u> N.L., C.H.; <u>Investigation:</u> N.L., C.H., R.M; <u>Formal analysis:</u> D.P., N.L., E.K.; <u>Writing – original draft:</u> N.L., E.K.; <u>Writing – review and editing:</u> B.H., E.K.; <u>Project administration:</u> E.K.

## Supplementary methods and results

**Table S1.** DIA variable windows used for Thermo Orbitrap HFX analysis.

| DIA Window | Centre Mass (m/z) | Start (m/z) | End (m/z) | Width (m/z) |
| --- | --- | --- | --- | --- |
| 1 | 372 | 350 | 394 | 44 |
| 2 | 408.5 | 393 | 424 | 31 |
| 3 | 437.5 | 423 | 452 | 29 |
| 4 | 464.5 | 451 | 478 | 27 |
| 5 | 490.5 | 477 | 504 | 27 |
| 6 | 516 | 503 | 529 | 26 |
| 7 | 541.5 | 528 | 555 | 27 |
| 8 | 567.5 | 554 | 581 | 27 |
| 9 | 594 | 580 | 608 | 28 |
| 10 | 621 | 607 | 635 | 28 |
| 11 | 648.5 | 634 | 663 | 29 |
| 12 | 677.5 | 662 | 693 | 31 |
| 13 | 708.5 | 692 | 725 | 33 |
| 14 | 741.5 | 724 | 759 | 35 |
| 15 | 778 | 758 | 798 | 40 |
| 16 | 819 | 797 | 841 | 44 |
| 17 | 866 | 840 | 892 | 52 |
| 18 | 925 | 891 | 959 | 68 |
| 19 | 1010 | 958 | 1062 | 104 |
| 20 | 1355.5 | 1061 | 1650 | 589 |

**Table S2.**
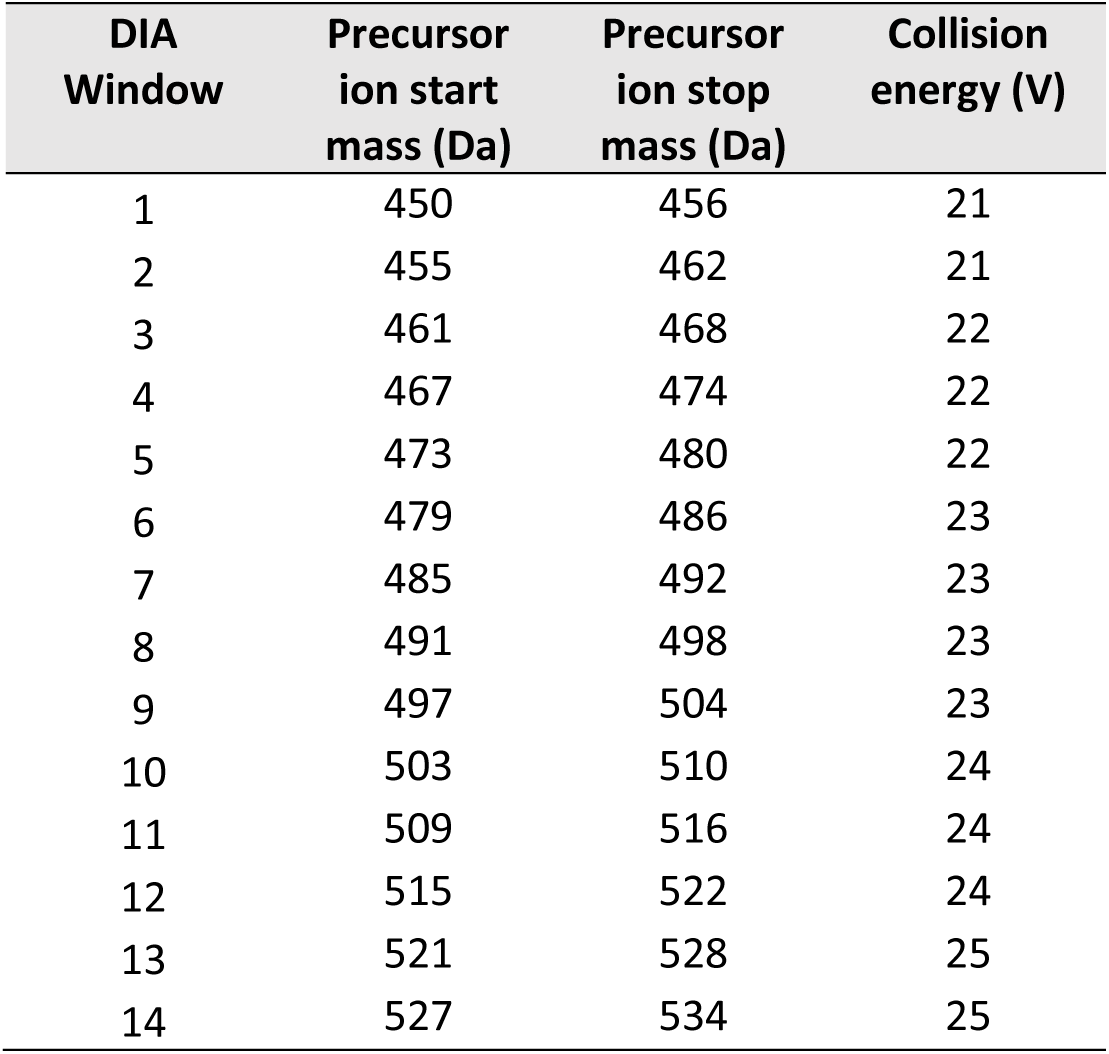
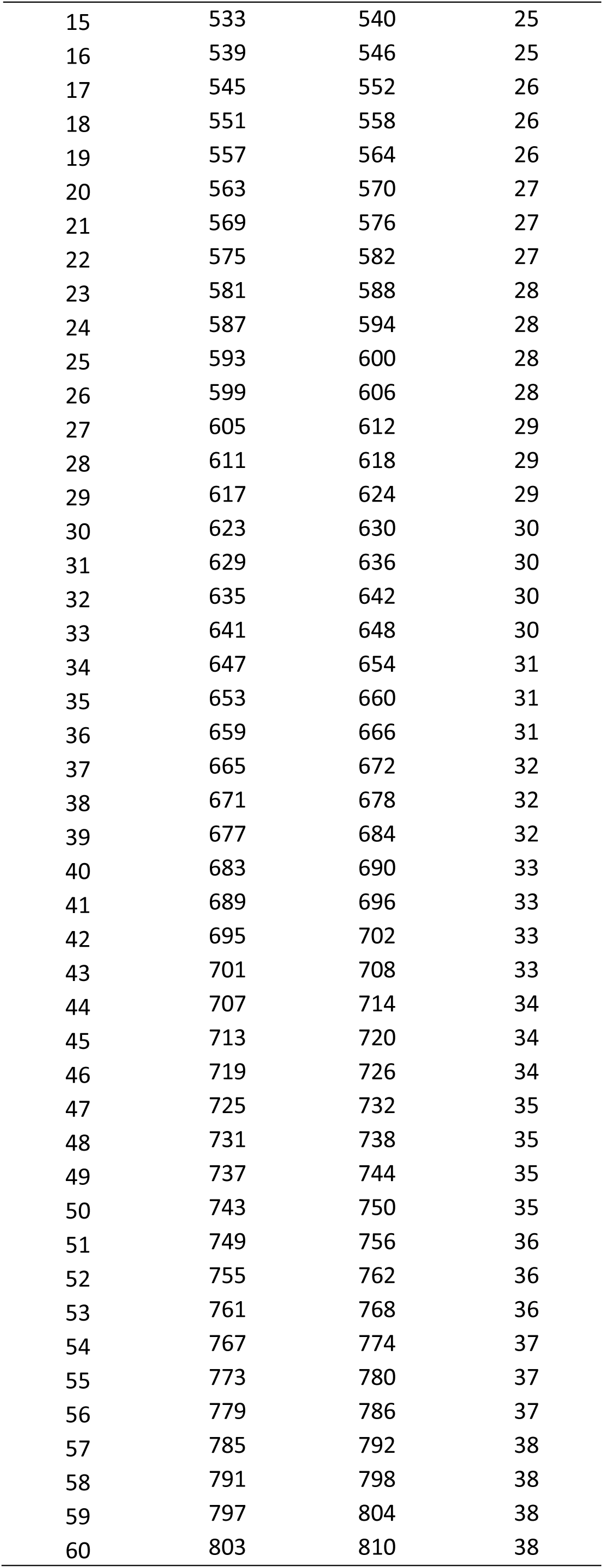
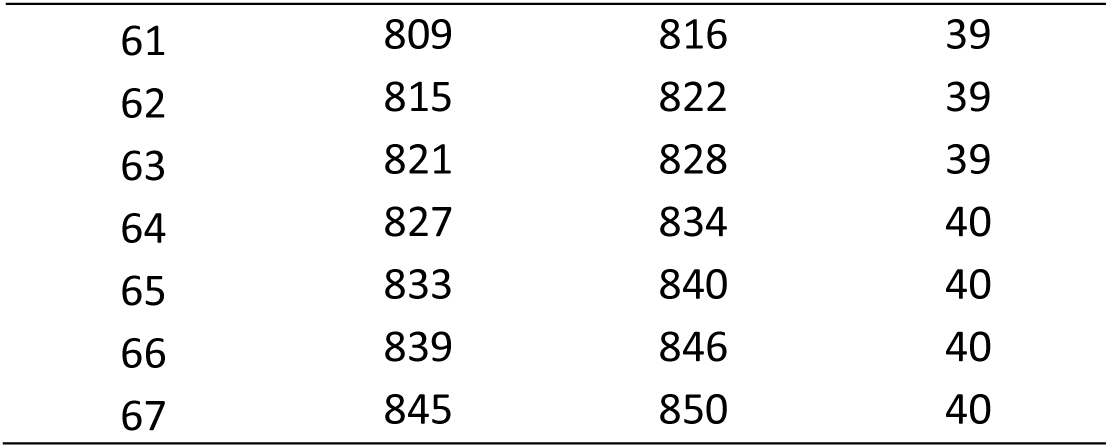
DIA variable windows used for ABSciex ZenoTOF 7600 analysis.

| DIA Window | Precursor ion start mass (Da) | Precursor ion stop mass (Da) | Collision energy (V) |
| --- | --- | --- | --- |
| 1 | 450 | 456 | 21 |
| 2 | 455 | 462 | 21 |
| 3 | 461 | 468 | 22 |
| 4 | 467 | 474 | 22 |
| 5 | 473 | 480 | 22 |
| 6 | 479 | 486 | 23 |
| 7 | 485 | 492 | 23 |
| 8 | 491 | 498 | 23 |
| 9 | 497 | 504 | 23 |
| 10 | 503 | 510 | 24 |
| 11 | 509 | 516 | 24 |
| 12 | 515 | 522 | 24 |
| 13 | 521 | 528 | 25 |
| 14 | 527 | 534 | 25 |
| 15 | 533 | 540 | 25 |
| 16 | 539 | 546 | 25 |
| 17 | 545 | 552 | 26 |
| 18 | 551 | 558 | 26 |
| 19 | 557 | 564 | 26 |
| 20 | 563 | 570 | 27 |
| 21 | 569 | 576 | 27 |
| 22 | 575 | 582 | 27 |
| 23 | 581 | 588 | 28 |
| 24 | 587 | 594 | 28 |
| 25 | 593 | 600 | 28 |
| 26 | 599 | 606 | 28 |
| 27 | 605 | 612 | 29 |
| 28 | 611 | 618 | 29 |
| 29 | 617 | 624 | 29 |
| 30 | 623 | 630 | 30 |
| 31 | 629 | 636 | 30 |
| 32 | 635 | 642 | 30 |
| 33 | 641 | 648 | 30 |
| 34 | 647 | 654 | 31 |
| 35 | 653 | 660 | 31 |
| 36 | 659 | 666 | 31 |
| 37 | 665 | 672 | 32 |
| 38 | 671 | 678 | 32 |
| 39 | 677 | 684 | 32 |
| 40 | 683 | 690 | 33 |
| 41 | 689 | 696 | 33 |
| 42 | 695 | 702 | 33 |
| 43 | 701 | 708 | 33 |
| 44 | 707 | 714 | 34 |
| 45 | 713 | 720 | 34 |
| 46 | 719 | 726 | 34 |
| 47 | 725 | 732 | 35 |
| 48 | 731 | 738 | 35 |
| 49 | 737 | 744 | 35 |
| 50 | 743 | 750 | 35 |
| 51 | 749 | 756 | 36 |
| 52 | 755 | 762 | 36 |
| 53 | 761 | 768 | 36 |
| 54 | 767 | 774 | 37 |
| 55 | 773 | 780 | 37 |
| 56 | 779 | 786 | 37 |
| 57 | 785 | 792 | 38 |
| 58 | 791 | 798 | 38 |
| 59 | 797 | 804 | 38 |
| 60 | 803 | 810 | 38 |
| 61 | 809 | 816 | 39 |
| 62 | 815 | 822 | 39 |
| 63 | 821 | 828 | 39 |
| 64 | 827 | 834 | 40 |
| 65 | 833 | 840 | 40 |
| 66 | 839 | 846 | 40 |
| 67 | 845 | 850 | 40 |

**Figure S1.**
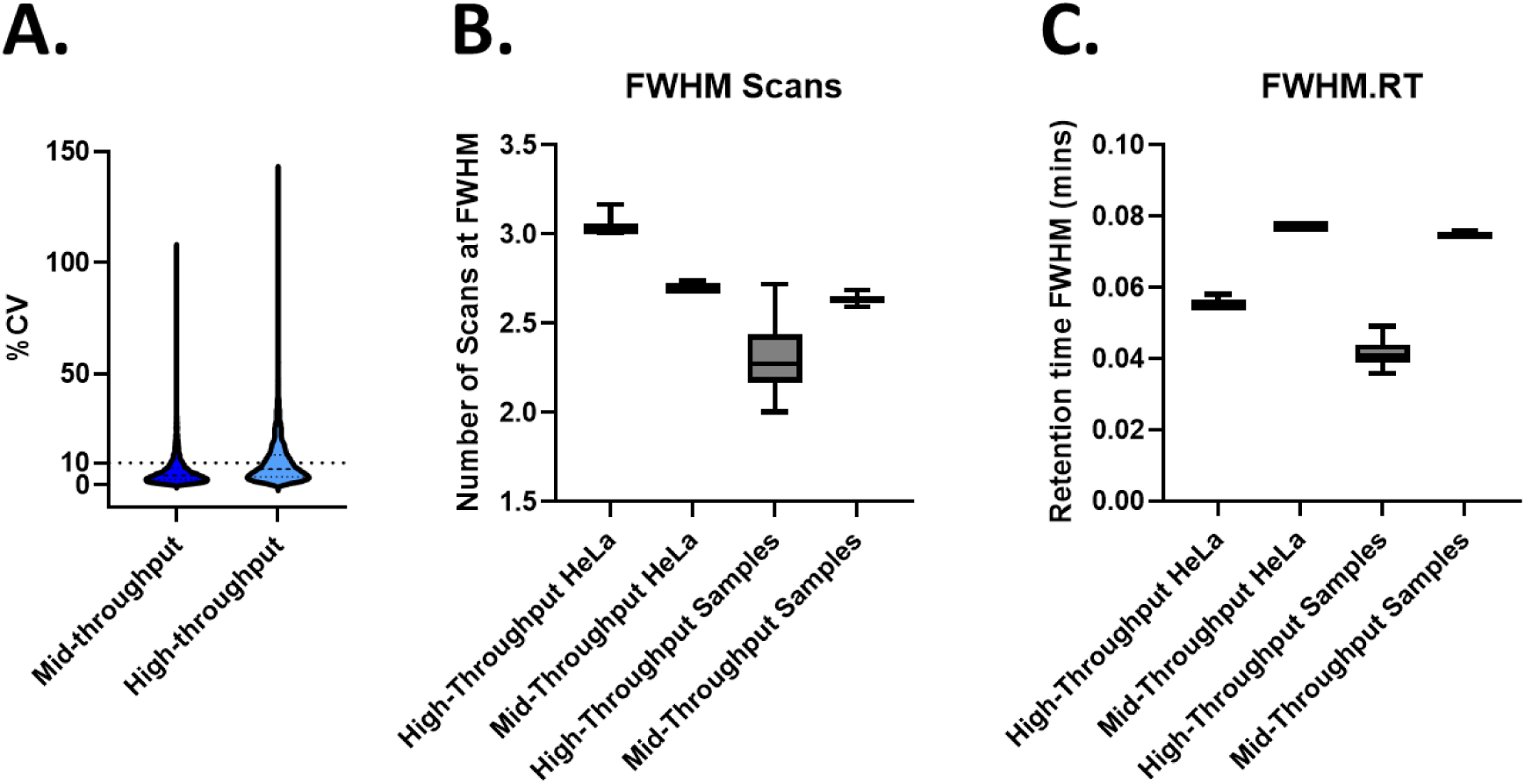
HeLa %CVs for proteins for both mid- and high-throughput methods (n=3) (A). Number of scans (B) and retention time (RT) (C) at FWHM for both HeLa (n=3) and samples (n=34) as reported via DIA-NN.

**Table S3.** List of differentially expressed proteins identified with both mid-throughput and high-throughput methods.

| Protein ID | Gene | Protein name |
| --- | --- | --- |
| Q9Y6N5 | SQOR | Sulfide:quinone oxidoreductase, mitochondrial |
| Q8IX19 | MCEMP1 | Mast cell-expressed membrane protein 1 |
| Q5QGZ9 | CLEC12A | C-type lectin domain family 12 member A |
| Q14739 | LBR | Delta(14)-sterol reductase LBR |
| Q13488 | TCIRG1 | V-type proton ATPase 116 kDa subunit a 3 |
| Q6UW68 | TMEM205 | Transmembrane protein 205 |
| Q10588 | BST1 | ADP-ribosyl cyclase/cyclic ADP-ribose hydrolase 2 |
| P62805 | H4C1 | Histone H4 |
| P33121 | ACSL1 | Long-chain-fatty-acid--CoA ligase 1 |
| P32942 | ICAM3 | Intercellular adhesion molecule 3 |
| P61626 | LYZ | Lysozyme C |
| P24158 | PRTN3 | Myeloblastin |
| P24071 | FCAR | Immunoglobulin alpha Fc receptor |
| P20292 | ALOX5AP | Arachidonate 5-lipoxygenase-activating protein |
| P20160 | AZU1 | Azurocidin |
| P18428 | LBP | Lipopolysaccharide-binding protein |
| P14780 | MMP9 | Matrix metalloproteinase-9 |
| P41218 | MNDA | Myeloid cell nuclear differentiation antigen |
| P11215 | ITGAM | Integrin alpha-M |
| P05164 | MPO | Myeloperoxidase |
| P05107 | ITGB2 | Integrin beta-2 |
| P04839 | CYBB | Cytochrome b-245 heavy chain |
| P02788 | LTF | Lactotransferrin |
| O95498 | VNN2 | Pantetheine hydrolase VNN2 |
| O75923 | DYSF | Dysferlin |
| P08670 | VIM | Vimentin |
| P08575 | PTPRC | Receptor-type tyrosine-protein phosphatase C |
| P08311 | CTSG | Cathepsin G |
| O75367 | MACROH2A1 | Core histone macro-H2A.1 |
| O00602 | FCN1 | Ficolin-1 |
| A8MVS5 | HIDE1 | Protein HIDE1 |

